# Diel and seasonal rhythmicity in activity and corticosterone in an Arctic migratory herbivore: A multifaceted approach

**DOI:** 10.1101/2024.08.30.610510

**Authors:** Margje E. de Jong, Annabel J. Slettenhaar, Rienk W. Fokkema, Marion Leh, Mo A. Verhoeven, Larry Griffin, Eva Millesi, Børge Moe, Elisabeth Barnreiter, Maarten J.J.E. Loonen, Isabella B.R. Scheiber

**Affiliations:** Dept. of Behavioral and Cognitive Biology, University of Vienna, Djerassiplatz 1, 1030 Vienna, Austria; Faculty of Biosciences and Aquaculture, Nord University, Kongens Gate 42, 7713 Steinkjer, Norway; Arctic Centre, Aweg 30, 9718 CW Groningen, The Netherlands; Conservation Ecology Group, Groningen Institute for Evolutionary Life Sciences (GELIFES), University of Groningen, Nijenborgh 7, 9747 AG, Groningen, The Netherlands; BirdEyes, Centre for Global Ecological Change at the Faculties of Science & Engineering and Campus Fryslân, University of Groningen, Wirdumerdijk 34, 8911 CE Leeuwarden, The Netherlands; NIOZ Royal Netherlands Institute for Sea Research, PO Box 59, 1790 AB Den Burg, The Netherlands; Department of Animal Ecology, Netherlands Institute of Ecology (NIOO-KNAW), Wageningen, the Netherlands; ECO-LG Ltd, Crooks House, Mabie, DG2 8EY, United Kingdom; Norwegian Institute for Nature Research (NINA), PO Box 5685 Torgarden, 7485 Trondheim, Norway

**Keywords:** Camera trap, GPS transmitter, circadian rhythmicity, hormones, continuous daylight, polar

## Abstract

Birds that migrate from temperate areas to the Arctic to breed lose their strongest *Zeitgeber* of circadian organization when they cross the Arctic circle in spring: the 24h light-dark cycle. Under continuous daylight, diverse behavioural and physiological patterns have been detected in both free-ranging and laboratory animals. To better understand the evolution of plasticity in circadian clocks, it is essential to study behavioural and physiological rhythmicity in the context of a species’ ecology. Employing a multifaceted approach, which included wildlife cameras, accelerometers, and non-invasive sampling of hormone metabolites, we investigated activity patterns and corticosterone rhythmicity in a migratory herbivore, the barnacle goose (*Branta leucopsis*), during its Arctic breeding season in Svalbard. We found that females showed a combination of both ultradian and diel rhythmicity in nest recesses and sleep during incubation. In both parents, these rhythms in activity continued also during the gosling rearing period. During moult, many geese aligned activity with the tidal rhythm. Barnacle geese showed weak diel rhythmicity in excreted corticosterone metabolites. This suggests that while Arctic geese may adopt an alternative *Zeitgeber* during the Arctic summer to maintain a diel rhythm, ultradian rhythmicity remains essential, allowing the geese to flexibly adjust their rhythms to environmental conditions.

## 1. Introduction

Various biological processes occur rhythmically, restricted to specific times of the day or season [1, 2], and in many species are regulated by photoperiod – the environmental period of daylight – as a *Zeitgeber* [3]. Upholding these rhythms in polar regions is challenging insofar as above the Arctic and below the Antarctic circle (66°N and S) natural light is completely absent in winter but continuously present during the summer [4]. Nevertheless, various Arctic species are able to retain endogenous and behavioural rhythms [reviewed in 1, 5].

Arctic vertebrates show variation in rhythmicity in behavioural activity between species, between individuals and within individuals. Arctic species are highly variable in their behavioural activity with some species (i) being completely *arrhythmic,* (ii) showing *circadian or diel rhythmicity, i.e*. displaying a recurring rhythm of approximately 24-hours even in the absence of light fluctuations, (iii) showing *ultradian rhythmicity* of recurrent periods of foraging and resting bouts less than 24-hours, or (iv) showing *free running rhythmicity, i.e*. not synchronized with environmental time cues [1, 5]. For herbivore species, Bloch *et al*. suggested that ultradian activity patterns in polar regions are linked with feeding behaviour and digestive processes [6]. Between individual differences exist, for example, in migratory shorebirds displaying bi-parental care, where conspecific pairs may show different incubation rhythms even when they breed in the same area [5, 7]. Furthermore, patterns may vary seasonally within individuals, *e.g.* driven by annual changes of day length [5, 6]. For example, the Svalbard rock ptarmigan (*Lagopus muta hyperborea*), the sole permanent resident bird on Svalbard, exhibits ultradian activity patterns with periodic feeding during polar summer and winter days. In contrast, during spring and fall its activity patterns are diurnal and feeding mainly occurs within daylight hours [8]. The diversity in behavioural responses to continuous daylight underlines the plasticity of the circadian system and indicates the relevance of the species’ biology [5, 9].

Behavioural and physiological processes are regulated through the rhythmic excretion of hormones by an endogenous network of one or several ‘Master’ clocks, which then coordinate and synchronise more peripheral clocks [reviewed in 1, 2]. Crucial for tracking environmental cycles is the neurohormone melatonin. Its diel secretion pattern, *i.e.* an extended peak at night and basal secretion during the day, provides the paramount hormonal signal transducing day length for peripheral clocks [3]. Melatonin secretion drives lasting changes in the hypothalamic-pituitary adrenal axis (HPA), where low levels of melatonin culminate in the increased secretion of the adrenal glucocorticoids, *i.e.* cortisol in mammals and corticosterone (CORT) in birds [1, 10, 11]. Glucocorticoids are (i) involved in correlating peripheral clocks [12], (ii) best known as being the downstream effectors of one of the two major stress response systems [13], and (iii) interrelate with feeding, modulation of energy storage and mobilisation as well as activity [10]. They display robust diel rhythmicity in latitudes with year-round day-night cycles [14-16, but see 17]. In temperate zones, the diel rhythmic pattern of baseline CORT correlates with activity; low levels of CORT during inactive phases are followed by a rise just before activity begins and CORT remains high as long as birds are active [10]. Contrary, in polar species with 24 hours of continuous natural light during the summer, findings of a diel CORT rhythm are ambiguous. For example, common murres (*Uria aalge*) maintain a diel CORT rhythm in the polar summer [18]. In contrast, the link between activity, time of day and corticosterone is not evident in thick-billed murres (*U. lomvia*) [10]. Other species without a diel CORT rhythm are Adélie penguins (*Pygoscelis adeliae*) [4] and common eiders (*Somateria mollissima*) [19]. Arrhythmicity in CORT was attributed to the suppression of melatonin secretion when photoperiod as *Zeitgeber* is absent [5, 10, 19]. Also, the lack of a rhythm in CORT in common eiders was linked to CORT interfering with the need to constantly forage [5], while in thick-billed murres it was ascribed to a stable modulation of energy storage and mobilisation [10].

Many bird species, which show rapid migration to their Arctic breeding grounds, experience large changes in daylight conditions en route, before they arrive in the polar regions with 24-hours of natural light. Latitudinal migrants may benefit from longer photoperiods at higher latitudes as for diurnal animals daily activity and foraging time can be increased [20]. Nevertheless, such quick fluctuations in light conditions present a significant challenge for circadian physiology [21–23]. There is a surprising variation in the response of migrants to continuous daylight [e.g. 5, 7], despite the fact that changing to a different rhythm may potentially be costly [24]. To increase our understanding of the evolution of plasticity in circadian clocks in migratory birds, it is thus necessary to study rhythmicity in behaviour and physiology in relation to the ecology of the species [e.g. 6, 7, 9].

Here we aim to quantify behaviour- and CORT-based rhythmicity in the barnacle goose, a migratory herbivore, during its breeding season in the Arctic using a multifaceted approach. Earlier findings on rhythmicity in barnacle goose behaviour and physiology are somewhat equivocal. A recent study showed that barnacle geese lost diel rhythmicity in body temperature once they encountered continuous daylight during their migration to the Arctic breeding sites [23]. During incubation in the Arctic, however, a diel rhythm in incubation recesses was described [25–27]. Likewise, young barnacle geese retained a diel, albeit time-shifted, pattern of corticosterone metabolites (CORTm) determined from droppings, over one 24-hour period in their first summer [28]. Here, we investigated potential diel and seasonal rhythmicity in activity and daily excretion patterns of CORTm in adult barnacle geese, using three different approaches; (i) activity and sleep during incubation, which were determined using photos from wildlife camera’s placed in the vicinity of the nest, (ii) activity across the breeding season of females and males, which was assessed using data from accelerometers (ACC), and (iii) CORTm concentration, which was determined from individually assigned droppings. Contrary to mammalian herbivores, where digestive processes dictate ultradian activity patterns [29, 30], barnacle geese do not possess complex digestive systems. However, their summer food retention time is increased by interrupting feeding with longer loafing spells [29]. We expected that barnacle geese would feed around the clock [*sensu* 23], which has been suggested to be a possible adaptation of Arctic herbivores to optimally utilize the peak in high quality food during the short summer [31]. If ultradian rhythmicity in activity is important, then we expected to find a weak or absent diel rhythm in CORTm. In contrast, if barnacle geese rely on an evolutionary-based endogenous clock to schedule their biology or respond directly to external diel cues, then we predicted diel rhythmicity in activity and CORTm throughout the breeding season, similar to some other polar species [e.g. 18, 32, 33].

## 2. Materials and Methods

### (a) Study site and species

We collected data during the summers of 2020 – 2022 in a barnacle goose population nesting on islets in Kongsfjorden, Svalbard. Geese from this population migrate annually from their wintering grounds in the United Kingdom to Svalbard, stopping along the Norwegian coast [29], and thereby experiencing a wide range of daylight conditions. Except on days with inclement weather when boating was not possible or polar bears (*Ursus maritimus)* were present, we monitored nests every other day during incubation and hatching on the two main breeding islands of the study area, Storholmen and Prins Heinrichøya from June to the beginning of July (see [34, 35] for details). We determined the exact nest location using a handheld GPS (Garmin GPSmap 64S), noted clutch size and identified parents by their individually recognizable engraved plastic leg rings when the researcher approached the nest [35]. To minimize disturbance, we got information on possible nests of geese with transmitters (see below) on other islands, *i.e.* Midtholmen, Juttaholmen, Observasjonsholmen, from colleagues who worked there regularly. Egg laying and incubation in barnacle geese takes approximately 29 days [36]. After hatching, many geese raise their goslings in or in the direct vicinity of the village Ny-Ålesund (78°55′30″N 11°55′20″E) [37, 38]. Here, we observed geese at least twice-daily using binoculars or spotting scopes to identify individuals by their colour rings, to establish family size and record feather moult. From the first day of moult, the complete moulting process lasts 35 to 40 days on average, with a flightless period of ca. 25 days [39]. In our analyses, we defined the flightless period to start at the first observation of loss of all flight feathers, and to end 25 days later (from now on called ‘moult’).

In 2020, we fitted 12 females and 12 males from 24 pairs with black coloured OrniTrack-NL40 3G transmitter neckbands (Ornitela, UAB, Lithuania. Mass ∼20 gram, height 21mm, diameter 38-40 mm). We caught ten geese on or near their nest on the islands Storholmen or Prins Heinrichøya during late incubation using two methods: (1) using a ∼ 6 m hand-held noose pole to catch the bird around its neck, or (2) by catching the geese by hand, which was mainly possible for aggressive males that came close. All remaining geese were caught during annually performed catches, during which flightless moulting geese are driven into corrals [29]. The 24 individuals were chosen following a suite of criteria, including that it would be desirable if both pair partners were marked with unique colour rings already (for details of criteria see [40]). We determined the sex of the birds from cloacal inspection [35]. During the incubation periods in 2021 and 2022, we deployed additional transmitters on three birds (one female and two males) and four birds (three females and one male), respectively.

### (b) Activity during incubation and across the breeding season

#### (i) Incubation patterns from wildlife camera photos

To investigate rhythmicity over the period of incubation, we set up wildlife cameras (Usogood TC30 Trail Camera) from 9/6/2021 to 16/7/2021 near the nests of incubating geese, which were fitted with GPS transmitters in 2020 (n = 15). Cameras, set in time-lapse mode, took two pictures every five minutes. We ultimately retrieved the cameras either after hatching or when a nest was preyed upon or abandoned [40].

We analysed pictures (n = 110739) with Timelapse2 Image Analyser (version 2.2.4.3, [41]). We quantified whether the female was (i) sitting on the nest, (ii) in ‘sleep posture’, *i.e.* head resting on back with beak often tucked under a wing [42], (iii) standing right next to the nest or (iv) absent from the nest. For further analyses of nest recesses, we pooled behaviours (iii) and (iv) as *active* and behaviours (i) and (ii) as *inactive*. For the analyses of ‘sleep’, we contrasted being in the sleep posture with the other behaviours. Males sometimes nest sit [40], but here we focussed on females only.

Instances, where males nest sat, while females were on incubation recesses, were classified as (iv) absent from nest. In total, wildlife cameras supplied data from 2 hours to 20 days. We only used data which exceeded the recommended minimum duration of ten days for biological rhythm analyses [43, 44]. We could assess activity, *i.e.* incubation recesses, and sleep patterns, from photos taken by the wildlife cameras in 11 out of 15 females. The other four cases were discarded, because data were insufficient due to nest (n = 2) or camera failure (n = 2).

#### (ii) Seasonal activity from transmitter accelerometer data

We used ACC data collected by the transmitters during the summers of 2021 and 2022 (2021: females n = 8, males n = 9; 2022: females n = 9, males n = 9; same individuals across 2021 and 2022: n = 12). The solar-powered transmitters were set to record a GPS-fix every 15 min when battery voltage was 75-100%, every 30 min at 50-74%, every 60 min at 25-49% or every 240 min at voltages lower than 25%. Immediately after each GPS-fix, a 2-sec ACC burst was taken at a frequency of 20 Hz. Gravitational acceleration was measured in unit g/1000 in the three spatial axes [45].

We used ACC data to identify activity and inactivity of individual geese during the breeding season [46, 47]. For each goose, we checked the number of measurements per burst taken after each GPS-fix to ensure a complete dataset. Then, we calculated the vectorial sum of dynamic body acceleration (VeDBA) from the ACC data, a common proxy for energy expenditure [48]. For this, we first calculated static acceleration, *i.e.* the average raw measured acceleration for each dimension (x, y, z) within the bursts. Second, we subtracted the static acceleration from the raw data for each dimension, thus getting dynamic acceleration. Last, we calculated the vectorial sum of dynamic body acceleration of each burst by taking the square root of the summed dynamic accelerations of each dimension [48]. Following [47], we created probability density histograms to identify peaks for activity and inactivity and used the *mix* function in the R-package *mixdist* to decompose the distribution into two gamma distribution components for active and inactive behaviour [49]. We found the threshold between the active and inactive behaviour distributions by calculating the intersection point between the two [see electronic supplementary material Figure S1, 46, 47].

In addition, we estimated the nesting period based on the method described by [45] using VeDBA and GPS data. Nesting is defined as the entire period of egg laying, nest building, incubation, and hatching. For females, we took daily median VeDBA < 1 for motionless days, as this corresponded best with observed periods of nesting in the field (n = 14: nesting period or part of the nesting period could be estimated, n = 3: nesting period could not be estimated due to transmitter attachment during hatching or when the goose was likely not breeding). We estimated the potential nest location by taking the median latitude and longitude of motionless fixes on days on which the goose was mostly motionless. Nest site attendance was calculated as the distance of each GPS-fix of a goose to its potential nesting location and then calculating the daily amount of time that the goose spent within a radius 50 m of its potential nest site. The attendance threshold was the first day on which the goose spent > 75% of time within 50 m of the nest and the period threshold was set at 3 days [45].

Most males, on the other hand, were never motionless during nesting (n = 14). Four males, however, did show motionlessness, which could plausibly be used to establish the nesting period. For nine additional males we established nesting based on field observations of the known nest location. For six males, however, neither method worked and we could not establish nesting.

In total, we obtained both an observed nest location in the field and an estimated nest location based on the transmitter data for 14 geese (males and females); the distance between the observed and estimated nest location was 2.37 meters on average (SD = 0.98, range 0.85 – 3.67; we excluded an outlier of 25.45 meters where the handheld GPS was likely turned on too late).

Later breeding stages, *i.e.* when geese had goslings and/or were moulting, were based on direct observations (see above). For 13 geese we obtained data on the period with goslings, for 11 geese when they were moulting and for three geese that had goslings while moulting (‘goslings & moult’ period). Since these different breeding stages pose different demands on the geese [29], we tested for rhythmicity within these three periods. Once again, we only used data, which exceeded the recommended minimum duration of ten days for biological rhythm analyses. Therefore, we excluded four nesting periods, five gosling periods, one gosling & moult period and one moult period, because here data collection time was too short. We pooled the remaining two goslings & moult periods with the moult period data to increase sample size.

### (c) Sample collection and corticosterone metabolite assay

To investigate diel rhythmicity of CORTm excretion, we collected droppings of ringed geese in the village of Ny-Ålesund from 1/7/2020 to 13/8/2020 (n = 349 droppings, 26 individuals; 14 females, 12 males) and 16/6/2021 to 31/7/2021 (n = 333 droppings, 52 individuals; 28 females, 24 males). For 17 individuals we collected samples in both years. We attempted to collect a minimum of three samples [50] per 3-hour time intervals, *i.e.* from 0:00 – 2:59, 3:00 – 5:59, …, 21:00 – 23:59 and intended to cover one complete 24-hour cycle every week for each individual. This was often not possible, as pairs sometimes left the area and could not be located. Only unambiguously assigned samples were collected and frozen at −20°C within 1 h (for methodology on how to collect droppings, see electronic supplementary material). Droppings were shipped frozen to the Dept. of Behavioural and Cognitive Biology, University of Vienna, Austria, for analyses.

Dropping samples for determining CORTm were quantified using an enzyme immunoassay validated for barnacle geese [51] and applied successfully in previous studies [*e.g.* 28]. CORTm was below the detection limit in three samples (total n = 679). Determined from homogenized pool samples, intra- and inter-assay coefficients of variation (% CV) were 11.91% (< 15%) and 13.03% (< 25%), respectively.

### (d) Statistical analyses

All analyses were performed in R version 4.4.0 (The R Foundation for Statistical Computing 2024).

#### (i) Rhythmicity in activity

We plotted actograms to visualize overall activity patterns during incubation, as assessed from wildlife camera data, and across the entire breeding season as determined from transmitter data (*ggetho* package [52]). Actograms, which show rhythmicity or a lack thereof, were double-plotted to facilitate inspection. We tested for periodicity in actograms using Lomb-Scargle periodogram (LSP) analysis [*periodogram* function, *zeitgebr* package: 52, 53] during specific periods: *i.e*. the limited time frame when cameras were placed during incubation until we spotted the first gosling on an image, and the entire nesting period (defined as the total time of nest building, egg laying, incubation, and hatching) as well as gosling rearing and moult for the transmitter data. The LSP analysis was suitable to discover periodicity in our dataset as it can handle unequally sampled time-series with missing data [53]. Periodicity, *i.e.* the regular recurrence of behaviour or an event over fixed, equal time intervals, is a common but not universal trait of rhythmicity, which in itself emphasizes the repetitive pattern of changes rather than precise timing [54]. For each dataset we ran two LSP analyses; one focussed on ultradian rhythmicity with a period range between 1 and 18 hours and another focused on diel rhythmicity with a period range between 18 and 36 hours [following 32, 44, 55]. Data were re-sampled every 5 minutes in the case of data from nest cameras and every 15 minutes for data provided by the transmitters. The oversampling rate was set at 100. We used the *find_peaks* function from the *zeitgebr* package to extract significant peaks (p < 0.05). Time-series data can contain two separate peaks, which may point to a true combination of ultradian and diel rhythmicity or it can be a data artefact [56]. We followed the methods of [55] to distinguish between these two: if we found two peaks in the normalized power for the same individual with an 18 h window we rejected the smaller peak if it was less than one-third the height of the larger one.

We investigated possible differences in ultradian and diel peak periods between breeding stages (nesting, goslings and moult) using linear mixed-effects models (*lme* from the package *nlme,*[57, 58]) with peak period in hours as the response variable and breeding stage (categorical) as a predictor variable. In addition, we also considered the predictor variables sex (categorical: female, male), the interaction between sex and breeding stage, and year (categorical). We accounted for repeated measures by adding ID as a random effect. We used automated model selection using the function *dredge* (package *MuMIn*) and AICc [59]. We give the 95% confidence set of models in the electronic supplementary material [60] and information including 85% confidence intervals for the full model in the text [61]. The 85% interval is consistent with how variables are selected using AIC. When present in the top model, we computed estimated marginal means for specific factors and comparisons among levels [package *emmeans*, 62].

#### (ii) Diel patterns of immune-reactive corticosterone metabolites

To investigate the association between predictor variables and CORTm concentration over the day, we fitted linear fixed-effects models (*lme*). We log-transformed CORTm concentration to adhere to model assumptions. The predictor variables in all models include time of day (continuous), year (categorical), day of the year (continuous) and sex (categorical). As time of day is a circular variable, we changed it into two linear variables by first transforming hour of day to radians and then calculating the sine and cosine of those radians. We included sine and cosine as continuous predictor variables in our models [18, 63]. To account for repeated measures of the same individuals, we added individual identity as random factor in all models. A more complex random slope model could not be explored, because this fitted mixed model was singular. As above, we used model selection using the function *dredge* (package *MuMIn*) and AICc and give the same model information.

## 3. Results

### (a) Rhythmicity in activity

#### (i) Rhythmicity during incubation

All females showed rhythmicity in incubation recesses (Figure 1A, for actograms of all females see electronic supplementary material Figure S2). Ten females showed a combination of both ultradian and diel rhythmicity while one female showed ultradian rhythmicity only (electronic supplementary material Table S1, Figure S3). The mean ultradian period in incubation recesses was 3.21 h (95% CI: 2.41, 3.89) and the mean diel period was 24.03 h (95% CI: 23.73, 24.38). Most incubation recesses occurred during the conventional day (Figure 1B). In addition, in nine females we found both ultradian and diel rhythmicity when resting in a sleep posture (Figure 1C, for actograms of all females see electronic supplementary material Figure S4), while two females showed diel rhythmicity only (electronic supplementary material Table S2, Figures S5). The mean peak ultradian period was 8.17 h (95% CI: 5.14,10.64) and the diel period was 24.96 h (95% CI: 23.57,1 26.12). Overall, geese were more often observed in the sleeping posture during conventional night time (Figure 1D).

**Figure 1:**
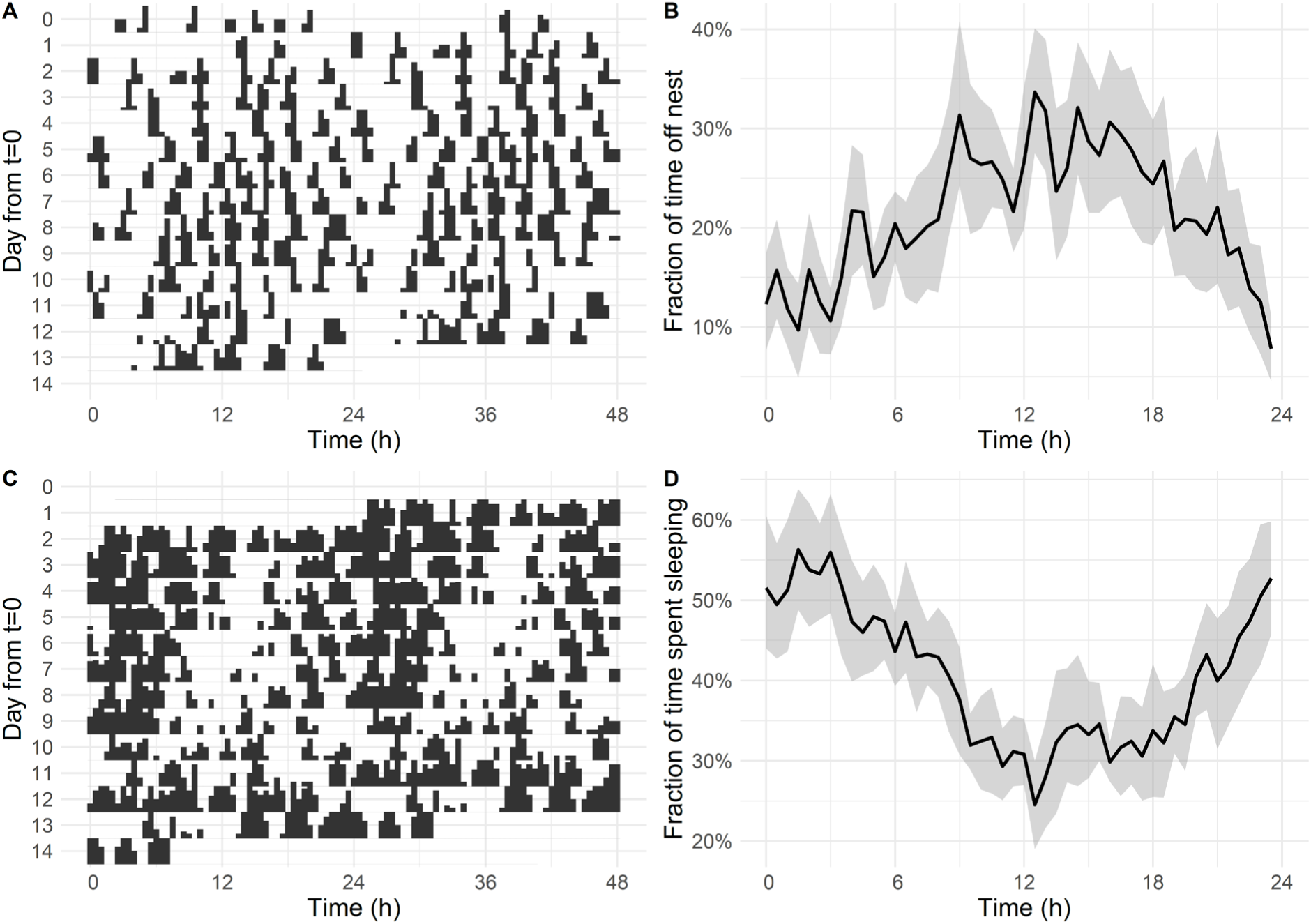
Rhythmicity of females during incubation recesses [panels (A) and (B)] and ‘sleep’ [panels (C) and (D)] based on wildlife camera pictures. Double plotted actograms of female ID CA41113 as an example show in black when she is away from the nest during incubation recesses (A) or when she is on the nest in the sleeping posture (C), while transparency indicates inactivity on the nest in (A) or any behaviour other than sleep in (C). The height of the bars indicates how often the specific behaviours are observed during that time. The x-axis displays two consecutive days and these consecutive days are also shown from top to bottom on the y-axis. On the y-axis t = 0 is the day the first pictures were taken by the wildlife cameras. Population level (N = 11) graphs show the fraction of time, averaged over 24 hours, spent on incubation recesses (B) and in sleep posture (D) in percentages (y-axis). The x-axis displays time over 24 hours, and the black line represents the mean, while in grey bootstrapped 95% confidence intervals are displayed.

#### (ii) Seasonal rhythmicity in activity

We detected rhythmicity in all geese over the course of the entire breeding season (for examples of actograms of two individuals see electronic supplementary material Figure S6). Except for three females, which showed no rhythmicity in activity patterns during nesting in 2022, most geese showed diel rhythmicity or a combination of ultradian and diel rhythmicity during nesting and when they had goslings (Figure 2 left and centre panel). Furthermore, we also detected either ultradian rhythmicity only or a combination of ultradian and diel rhythmicity during moult (Figure 2 right panel). We found evidence that breeding stage correlated with ultradian peak period, as this variable was present in all models within the 95% confidence set (Table 1A, electronic supplementary material Table S3). The top model included breeding stage only (Table 1A) and post-hoc testing revealed that the ultradian peak period during moult was significantly higher than when geese nested or had goslings (Figure 3, goslings – moult; estimate = −6.05, SE = 1.75, p = 0.015, goslings – nesting; estimate = 0.35, SE = 1.80, p = 0.9794, moult – nesting; estimate = 6.40, SE = 1.49, p = 0.004). There was no strong indication that breeding stage influenced diel peak period, with the top model being the null model (Table 1B, Figure 3, see also electronic supplementary material Table S4). The fraction of time when the geese were active, varied between males and females during nesting, with females being more active around noon, while males were less active then (Figure 4A). When geese had goslings or were moulting, females and males did not differ in the fraction of time they were active (Figure 4B & C), but there was a sharp decrease in activity after midnight until early morning. This pattern was not present during moult.

**Figure 2:**
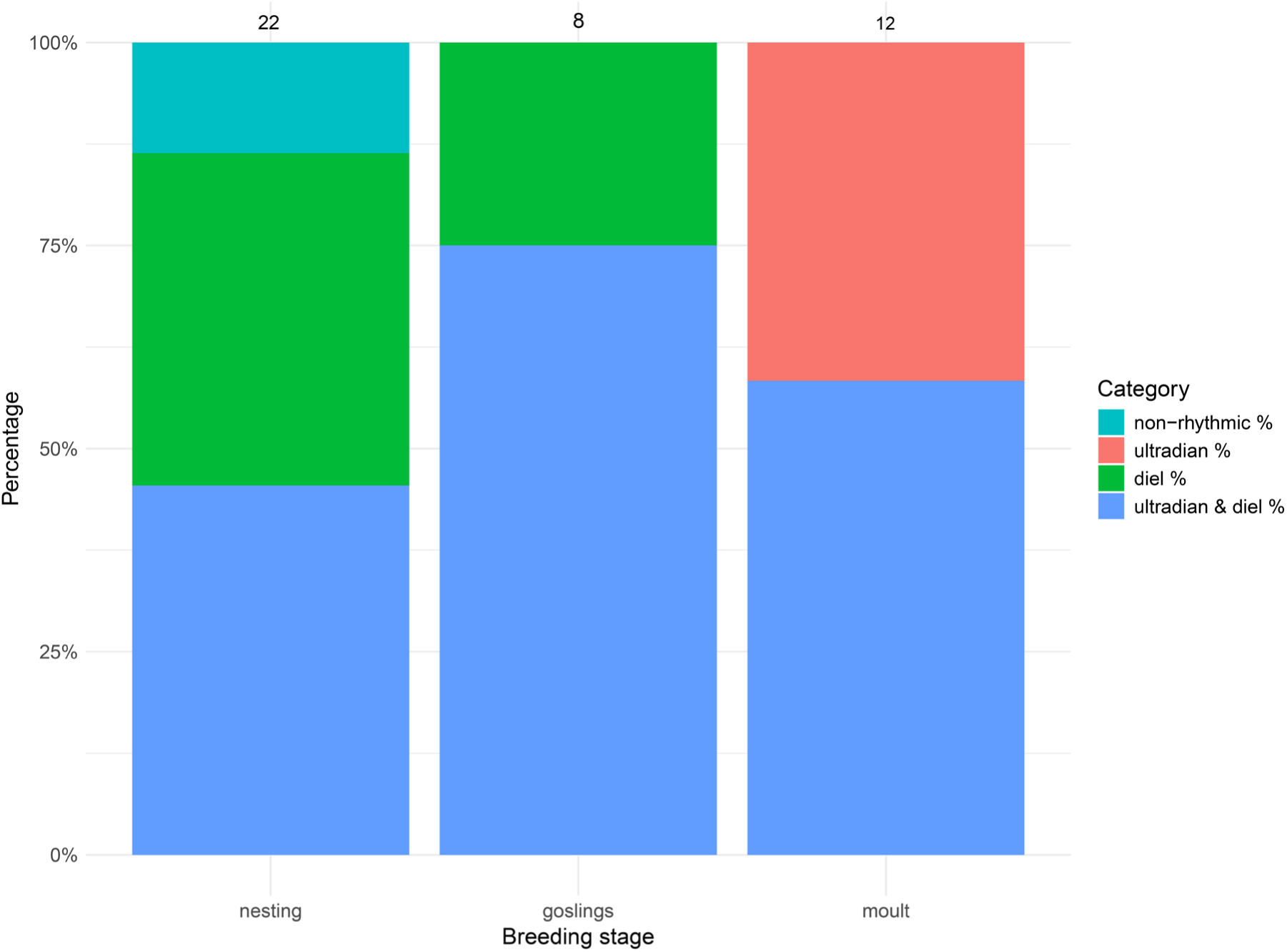
Percentages of rhythmicity types based on transmitter accelerometer data for 18 individual geese over the course of two summer seasons. Over the three breeding stages either ultradian or diel rhythmicity alone, a combination of ultradian and diel rhythmicity, or no rhythm were observed. Numbers above the stacked bars indicate sample size per breeding stage.

**Figure 3:**
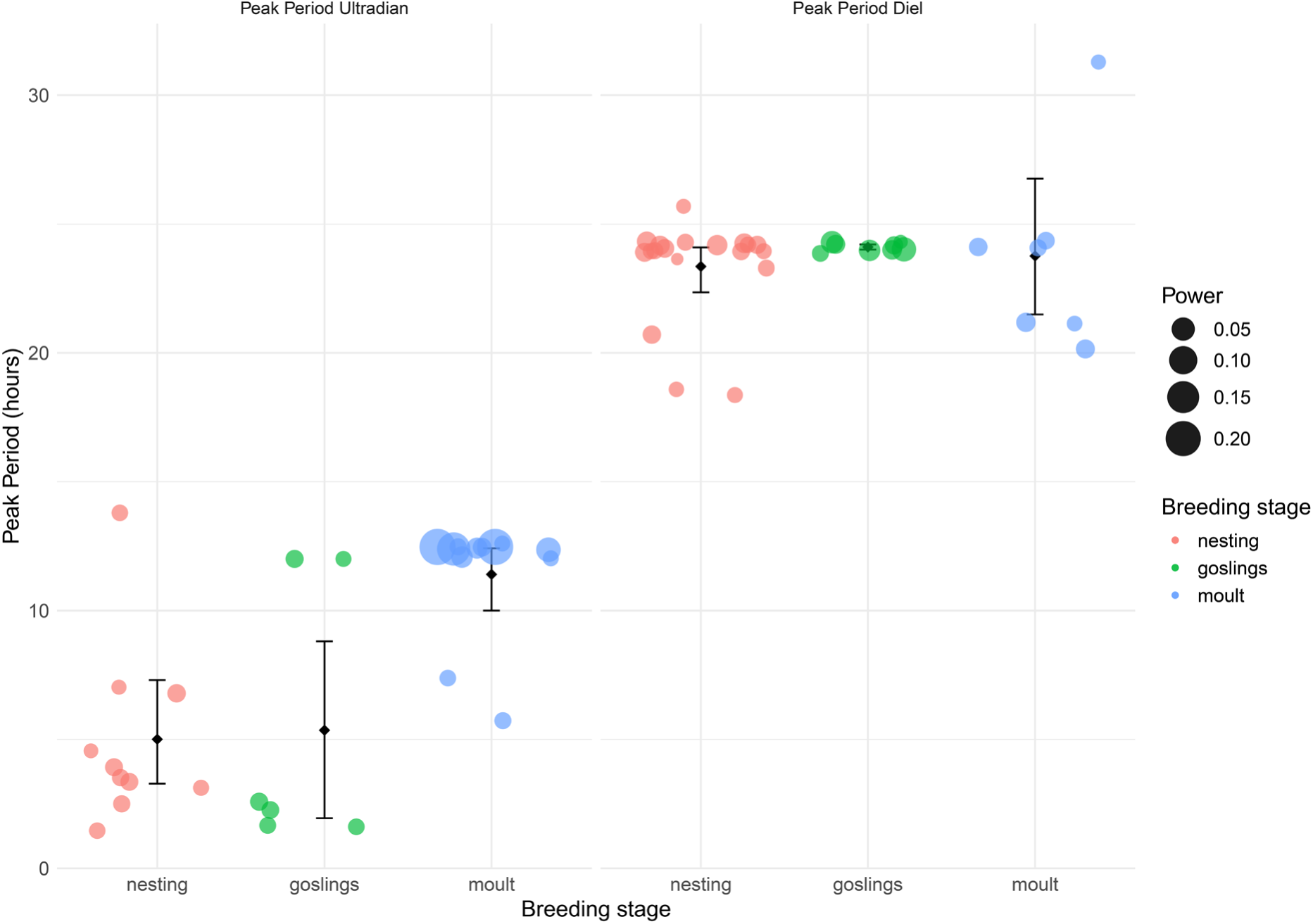
Diel and ultradian average peak periodicity during different breeding stages based on transmitter accelerometer data. The values of the peak period are for rhythmic geese only, i.e. a peak above the significance threshold at α = 0.05. Individual geese are shown as dots. The size of the dots represents the power of the peak period, i.e. a representation of the strength of the signal. The error bars are 95% bootstrap confidence intervals on the population mean.

**Figure 4:**
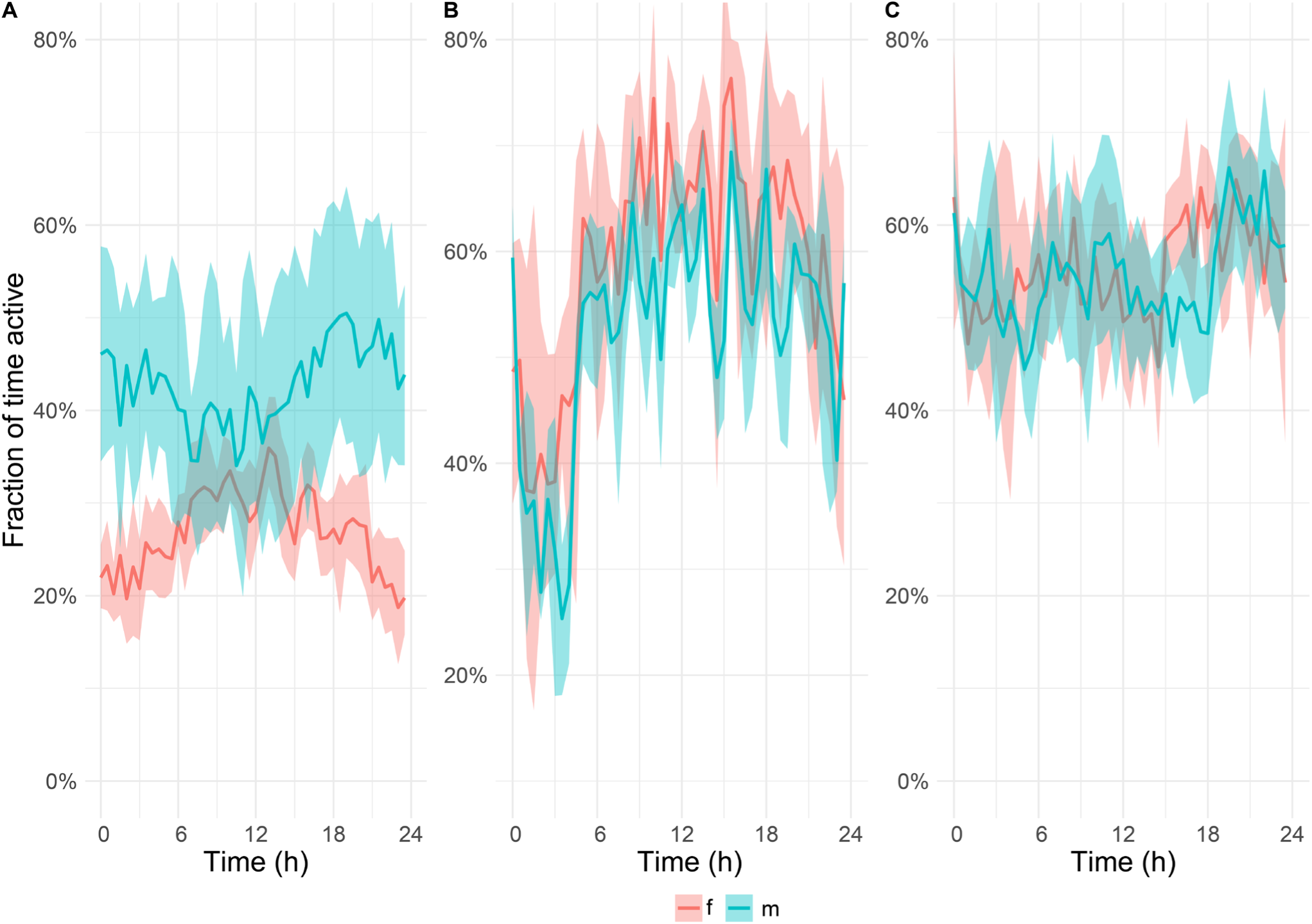
Population level graphs, based on transmitter accelerometer data, showing the fraction of time active, averaged over 24 hrs., during nesting (A), when geese had goslings (B), and during moult (C) in percentages (y-axis). Females (f) and males (m) are indicated in different colours. The x-axis displays time over 24 hours, and the solid lines represent the mean, while in transparent colour bootstrapped 95% confidence intervals are displayed.

**Table 1:**
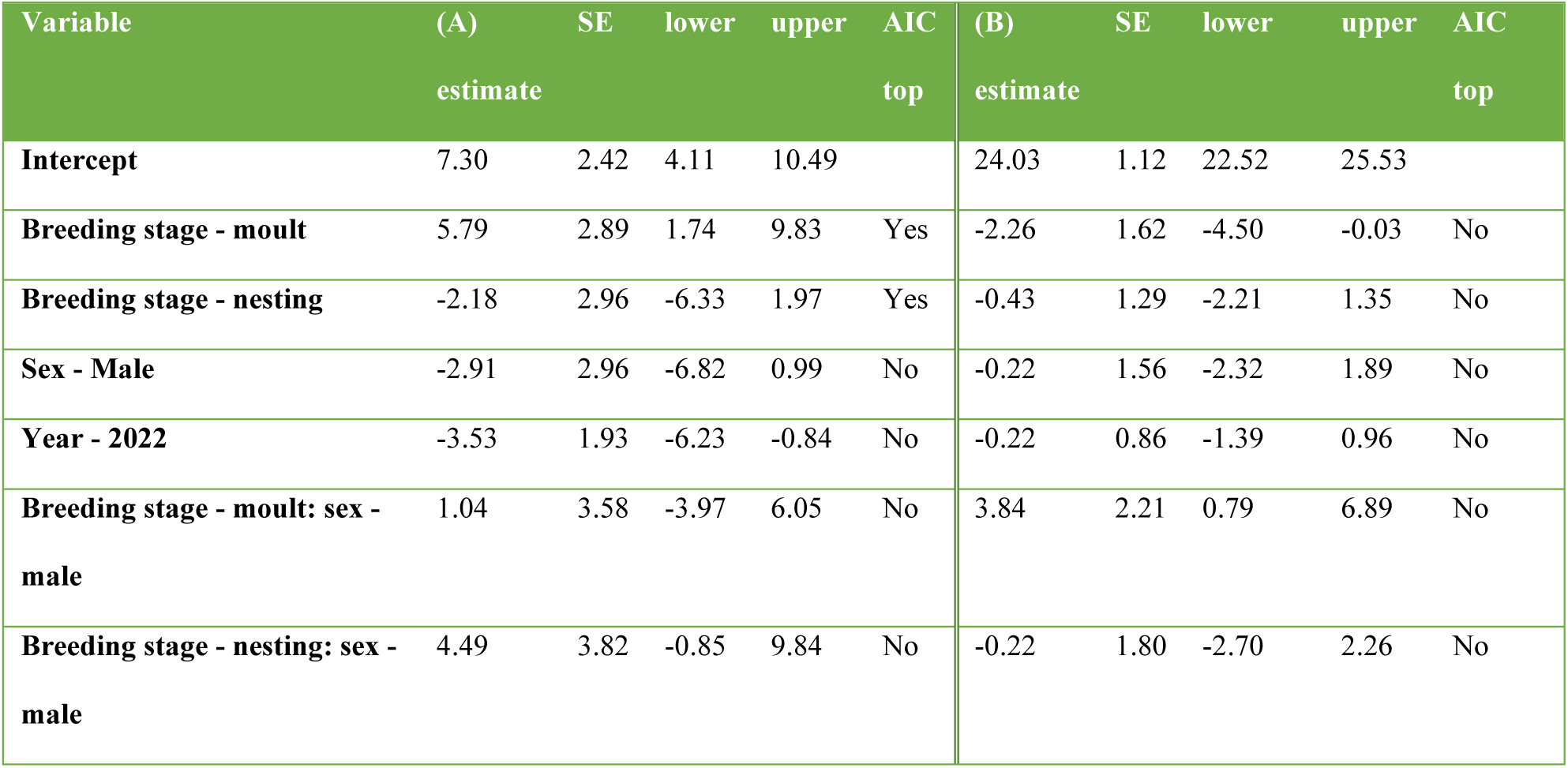
Intercept and coefficient estimates from the full models investigating differences in (A) ultradian peak period in hours and (B) diel peak period in hours with corresponding standard errors (SE), 85% Confidence intervals (lower, upper), and if the variable was selected in the top AIC model. For (A) the random intercept for individual ID had a standard deviation of 0.0001 and the residual standard deviation was 2.96. For (B) the random intercept for individual ID had a standard deviation of 0.87 and the residual standard deviation was 1.76.

| Variable | (A)<br>estimate | SE | lower | upper | AIC<br>top | (B)<br>estimate | SE | lower | upper | AIC<br>top |
| --- | --- | --- | --- | --- | --- | --- | --- | --- | --- | --- |
| **Intercept** | 7.30 | 2.42 | 4.11 | 10.49 |  | 24.03 | 1.12 | 22.52 | 25.53 |  |
| **Breeding stage - moult** | 5.79 | 2.89 | 1.74 | 9.83 | Yes | -2.26 | 1.62 | -4.50 | -0.03 | No |
| **Breeding stage - nesting** | -2.18 | 2.96 | -6.33 | 1.97 | Yes | -0.43 | 1.29 | -2.21 | 1.35 | No |
| **Sex - Male** | -2.91 | 2.96 | -6.82 | 0.99 | No | -0.22 | 1.56 | -2.32 | 1.89 | No |
| **Year - 2022** | -3.53 | 1.93 | -6.23 | -0.84 | No | -0.22 | 0.86 | -1.39 | 0.96 | No |
| **Breeding stage - moult: sex - male** | 1.04 | 3.58 | -3.97 | 6.05 | No | 3.84 | 2.21 | 0.79 | 6.89 | No |
| **Breeding stage - nesting: sex - male** | 4.49 | 3.82 | -0.85 | 9.84 | No | -0.22 | 1.80 | -2.70 | 2.26 | No |

### (b) Diel patterns of immune-reactive corticosterone metabolites (CORTm)

Our results revealed that CORTm concentration was marginally influenced by time of day (Table 2), with both cosine and sine of time of day being present in the top model as well as in many models within the 95% confidence set (electronic supplementary material Table S5). CORTm concentration increased during conventional night time and decreased during day time hours (Figure 5). CORTm concentrations were slightly different between the years (Table 2; 2020 median: 32.13 ng/g dropping, range: 7.49 – 320.9 ng/g; 2021 median: 28.56 ng/g dropping, range: 1.02 – 583.56 ng/g). In addition, over the course of the season CORTm concentration slowly declined (Table 2). Collected mostly late at night on one specific day, several samples contained very high levels of CORTm, *i.e.* >200 ng CORTm/g droppings (n = 7). Omitting these data points in a second analysis revealed that the outcome did not differ substantially, except that the top model now only included the cosine of time of day and year (electronic supplementary material Tables S6, S7).

**Figure 5:**
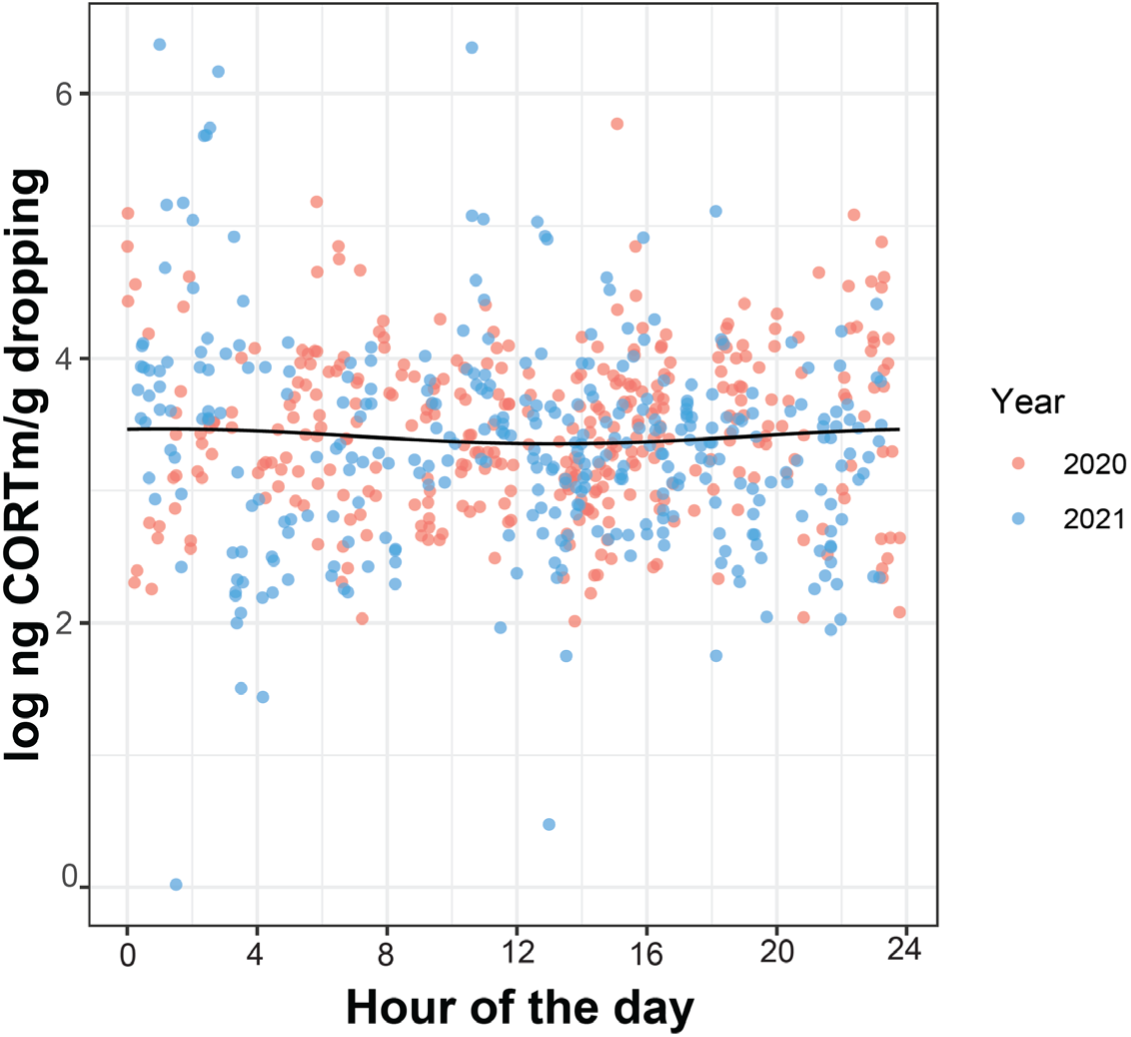
Daily rhythmicity in log-transformed corticosterone metabolite (CORTm) concentration measured in barnacle goose droppings in two years. The dots represent individual measurements and the solid line represents the predicted values from the model. 2020: n = 349 droppings, 26 individuals. 2021: n = 333 droppings, 52 individuals.

**Table 2:** Intercept and coefficient estimates from the full model investigating rhythmicity in log-transformed corticosterone metabolite concentration with corresponding 85% CIs, and if the variable was selected in the top AIC model. The random intercept for individual ID had a standard deviation of 0.23 and the residual standard deviation was 0.67.

| Variable | estimate | standard<br>error | lower | upper | AIC top |
| --- | --- | --- | --- | --- | --- |
| **Intercept** | 5.03 | 0.45 | 4.38 | 5.67 |  |
| **cosine** | 0.1 | 0.04 | 0.05 | 0.15 | Yes |
| **sine** | 0.06 | 0.04 | 0.003 | 0.11 | Yes |
| **sex - male** | -0.08 | 0.09 | -0.21 | 0.05 | No |
| **year - 2021** | -0.2 | 0.06 | -0.29 | -0.11 | Yes |
| **year day** | -0.007 | 0.002 | -0.01 | -0.004 | Yes |

## 4. Discussion

Svalbard barnacle geese maintain rhythmicity in behaviour and physiology during their Arctic breeding season. We detected that most females showed a combination of both ultradian and diel rhythmicity in incubation recesses and sitting in a sleep posture on the nest. Most females and males also showed a combination of ultradian and diel rhythmicity or diel rhythmicity only in activity during their entire nesting period or when they had goslings. During moult, geese exhibited either ultradian rhythmicity alone or a combination of ultradian and diel rhythmicity in their activity, possibly indicating a response to tidal rhythms (see below). We detected no differences in peak periodicity between females and males across the various breeding stages. However, during the nesting phase, females and males exhibited contrasting activity patterns, with females leaving their nest more often during the day and males being more active during the conventional night. Furthermore, the geese showed weak diel rhythmicity in excreted corticosterone metabolites with concentrations increasing during the night and decreasing during the day. Below we discuss our findings in more detail.

### (a) Ultradian rhythmicity in goose activity behaviour

Our results indicate the significance of ultradian rhythmicity, *i.e.* periods of less than 24 hours, in activity behaviour over the breeding season. During nesting and when geese had goslings, they showed ultradian activity rhythms with a period of ∼ 5 h (Figure 3). This ultradian rhythm might display a foraging activity – resting pattern [27], as is also shown by other Arctic herbivores such as Svalbard ptarmigan, Svalbard reindeer (*Rangifer tarandus platyrhynchus*) and muskoxen (*Ovibos moschatus*) during the summer solstice [e.g. 44, 55, 64, 65]. Such an activity pattern could be endogenously generated by ultradian oscillators [65] or controlled by the interplay of feeding and digestion [6, 66].

### (b) Diel rhythmicity in goose activity behaviour

Many geese also showed diel activity patterns in behaviour, *i.e.* periods of around 24 hours, throughout the breeding season. The presence of a diel rhythm does not inherently imply, however, that circadian systems are functional and/or entrained as the geese could be reacting directly to external cues, also known as “masking” [1, 67]. For example, during nesting, female geese could respond adaptively to temperature differences between conventional day and night, with slightly higher temperatures during the day despite 24h daylight. This might limit egg cooling rates during incubation recesses, as shown in snow geese [*Anser caerulescens*, 68]. Males do not incubate, although they sometimes sit on the nest [40], but often stand guard close by the nest when the female is on incubation recesses [27]. Nests, however, are more vulnerable to predation if the female is on recess [69] with the most significant egg predators, glaucous gulls (*Larus hyperboreus*) and Arctic skuas (*Stercorarius parasiticus*), being more active during conventional daytime [25]. This conflicts with females being on recesses then. Arctic foxes (*Aleopex lagopus*), on the other hand, are usually more active at night [25, pers. obs.]. For more than a decade, however, no foxes have been observed on those islands, where nests were monitored (pers. obs.). By contrast, foxes have been present in the gosling rearing areas during the study years. Here, males and females usually rested close to water bodies, into which they could escape from the fox [38, pers. obs.]. Although barnacle geese may not be able to protect their nests against foxes even if both parents are present [34, 70], fox predation was suggested to be the selective force of the observed diel rhythmicity during incubation [25] as a ‘ghost of competition past’. If the day-active behaviour of female geese during nesting is shaped by an expected daily rhythm in fox predation risk, it suggests the activity pattern is regulated by an endogenous, innate circadian mechanism [71]. It is feasible that a goose’s circadian clock continues to function by employing alternative potential *Zeitgeber*s, such as diurnal changes in light intensity, polarization patterns, solar azimuth, UV radiation, changes in the spectral composition of light or slight changes in ambient temperature, which exist under polar summer conditions [72].

### (c) Ultradian and diel rhythmicity in sleep

Studies on sleep patterns under natural conditions are rare due to challenges in measuring sleep in wild free-moving animals [73]. During incubation, we scored sleep posture, which has been associated with rapid eye movement (REM) sleep and non-REM (NREM) sleep in domesticated geese [42]. We therefore assume that our behavioural scoring of sleep was valid overall, although misclassifications may have occurred if individuals fell asleep while sitting with their beaks pointing forwards or were awake in the typical sleep posture. In addition, various bird species, such as mallards (*Anas platyrhynchos*) for example, perform unihemispheric sleep [74], which we could not measure here. The diel sleep pattern contrasts distinctly with the diel pattern observed in nest recesses (Figure 1), with females sleeping more during the conventional night. In addition, we detected ultradian rhythmicity, with a mean peak period of ∼ 8 h. Awakening bouts could serve as a periodic screening of the environment for danger [74, 75]. Captive barnacle geese kept under a natural day - night cycle also showed scattered sleep over the course of the day during summer [76], and the authors proposed that barnacle geese might have an attenuated circadian organization and may profit from becoming arrhythmic during the polar summer. However, we found no strong evidence for arrhythmia.

### (d) Tidal rhythm in goose activity during wing moult

During wing moult, barnacle geese are most vulnerable to fox predation [69]. At times, when goslings are still present, parents are forced to forage in areas with high quality food even during moult, thereby risking higher predation [38]. When goslings are preyed upon, adults can retreat to areas with lower-quality food but higher safety, such as tundra lake shores with mossy vegetation [38]. Over the past years, we noticed moulting geese without young to increasingly utilize the intertidal area to rest and/or forage on algae (pers. obs.). The ∼ 12 h ultradian rhythmicity patterns of geese during moult reflect the falling and rising tides (Figure 3, electronic supplementary material Figure S6A, [77]). The tides pose a constraint on an all-day-activity [78, 79], but the nearby sea provides safety from foxes. Additionally, the enhanced digestion during moult may help geese process and extract nutrients from this otherwise difficult-to-digest food [30, 80]. Although barnacle geese were shown to avoid salt-marsh vegetation in spring when it was experimentally sprayed with seawater, because they presumably are less capable of physiologically coping with very high salt loads in their environment [81], moulting geese foraged on algae in our study, possibly benefitting from fresh river water nearby. Moult in geese is energetically costly, because they moult wing coverts, some body feathers and flight feathers simultaneously. To compensate, geese become less active overall (Figure 4C, [39]). In contrast, geese, which still lead young, seem to retain their activity rhythm also throughout moult. This should be investigated in the future, because here sample sizes of geese with goslings were too small to confirm this finding.

### (e) Weak diel rhythmicity in corticosterone metabolites (CORTm)

Besides rhythmicity in activity, geese also maintained an, albeit weak, rhythmic pattern of excreted CORTm (Figure 5), similar to what we found in human-raised barnacle goslings over 24 hours [28]. Glucocorticoids are integral in providing physiological signals to regulate biological rhythms in sync with daily environmental cycles linking with activity and feeding [11, 12]. During the gosling period, when we collected the majority of samples for CORTm measurements, our finding contradicts this general assumption, because values dropped from conventional midnight to being lowest around midday, when geese are most active. As discussed in an earlier study [28] as well as above, increased fox activity at night, during the period when the goose families preferentially rest, could have affected the CORTm rhythm. In sum, whether the attenuated rhythm in CORTm is sufficient to regulate activity patterns and other functions during the polar summer or whether it plays only a minor role in endogenous timekeeping [18] needs to be investigated in the future.

### (f) Suitability of the methods used

Our use of a multifaceted approach, combining behavioural observations via wildlife cameras and accelerometers with corticosterone measurements, allowed for a comprehensive understanding of rhythmicity in barnacle geese during their Arctic breeding season in 24h daylight. We found accelerometers to be a suitable alternative for investigating activity, where time consuming direct behavioural observations are difficult. They are a valid substitute also for pictures obtained from wildlife cameras, where a multiplicity of photos need to be analysed in detail afterwards. For example, we found that the peak period in activity of five females during incubation, identified from both wildlife cameras and accelerometers corresponded very well. Except for one female, where we found diel rhythmicity retrieved from the ACC data which was not detected in data from the wildlife cameras, the ACC-based activity patterns gave reliable results in the other four individuals, despite the slightly longer 15 min intervals of ACC measurements relative to the 5 min intervals of subsequent photos (ultradian: camera data; mean = 3.5 h, SD = 1.7, ACC data; mean = 3.8 h, SD = 1.7. diel: camera data; mean = 23.8 h, SD = 0.7, ACC data; mean = 24.2 h, SD = 0.1). This indicates that ACC-based data described activity very reliably. This is supported by another study in captive barnacle geese, which investigated sleep – wake rhythms. Here, activity measurements from ACC data correlated well with patterns retrieved from electroencephalograms (EEG) [76]. Contrary to our study, where ACCs were mounted in neck collars, these ACCs were head-mounted. They took high-frequency measurements at a high sampling rate of 100 Hz, and could, thus, record even small head movements. In our study data were less fine-tuned, because they were collected at a lower rate, *i.e.* every 15 min during 2-sec bursts of 20 HZ. Therefore, we cannot provide the same inferences at present.

### (g) Conclusion

To summarize, over the course of their Arctic breeding season, barnacle geese show some degree of plasticity in their daily rhythms. Such intra-individual shifts in rhythms between sexes and breeding stages also exist in shorebirds [reviewed in 1, 5]. One possible explanation is that in barnacle geese the circadian clock mechanism keeps ticking, but the control, which it exerts over behavioural output, is plastic and is applied only when it provides some advantage [1, 2]. Alternatively, the observed rhythms are not endogenously controlled and geese respond directly to environmental or social cues, such as temperature, predation pressure, tides and/or their mates’ or goslings’ activity. The investigation of individual plasticity and consistency in behavioural and hormonal rhythmicity, and their relationship with fitness, are novel avenues for future work.

## Ethics

The study conforms to Directive 2010/63/EU and was conducted under FOTS ID 23358 from the Norwegian Animal Research Authority and approved by the Governor of Svalbard (RIS ID 11237).

## Data accessibility

Data and code used to conduct the analyses will be openly available in the Dryad data repository when the manuscript has been accepted for publication. Supplementary material will be available online.

## Declaration of AI use

As most authors are non-native English speakers, AI was occasionally used while writing to improve language and readability. AI was sometimes used as ‘search engine’ to find suitable code for graphics.

## Authors’ contributions

M.E.d.J.: Conceptualization, Data curation, Formal analysis, Investigation, Methodology, Project administration, Resources, Supervision, Validation, Visualization, Writing – original draft, Writing – review & editing; A.J.S.: Data curation, Formal analysis, Funding acquisition, Investigation, Methodology, Validation, Writing – review & editing; R.W.F.: Investigation, Methodology, Validation, Writing – review & editing; M.L.: Investigation, Validation; M.A.V.: Investigation, Writing – review & editing; L.G.: Methodology, Resources, Writing – review & editing; E.M.: Resources, Writing – review & editing; B.M.: Funding acquisition, Writing – review & editing; E.B.: Investigation, Validation, Supervision; M.J.J.E.L: Investigation, Resources, Supervision; I.B.R.S.: Conceptualization, Funding acquisition, Investigation, Methodology, Project administration, Resources, Supervision, Validation, Writing – original draft, Writing – review & editing. All authors gave final approval for publication and agreed to be held accountable for the work performed therein.

## Acknowledgments.

We are grateful to the staff of the French - German Arctic Research Base at Ny-Ålesund, chiefly Bettina Haupt (station leader AWI 2020/21), and KingsBay AS, Ny-Ålesund for logistic support, as well as Mindaugas Dagys (Ornitela – Ornitology and Telemetry Applications, Lithuania) for his helpful advice on transmitter technology. We thank everyone who observed geese in Ny-Ålesund in 2021 and 2022; Mikael Sætre, Simen Karlsen, Christophe Brochard, Anne Vorenkamp and Pyter Bootsma. We thank Kjetil Sagerup, Geir Wing Gabrielsen and Kjell Tore Hansen for sharing sightings of geese with GPS transmitters in Kongsfjorden, and Kees Schreven and Chiel Boom for R coding advice. Dropping samples were shipped with permission from the Federal Ministry Republic of Austria – Social Affairs, Health, Care and Consumer Production (Bescheide Nr. 2020-0.233.496, 2021-0256.249 – BMSGPK-Gesundheit – IX/B/10).

