## Supplement for "Diel and seasonal rhythmicity in activity and corticosterone in an Arctic migratory herbivore: A multifaceted approach"

### Supplement: Material and methods

##### *Dropping collection*

Individual droppings for determination of hormone metabolites were collected by following individual geese from a distance with a telescope. The scope was fixed on the exact spot at which a goose defecated. As soon as geese voluntarily left the area, the observer was guided to this spot by a second person with handheld VHF radios. It was relatively easy to locate the correct droppings, because of the shortly grazed tundra vegetation with small rocks, flowers *etc.* as reference points, and because new droppings are still wet relative to other dried up droppings in the vicinity. In cases of uncertainty to assign a dropping correctly, it was left behind.

### Supplement: Tables and Figures

##### *Incubation behaviour*

*Table 1: Results of the Lomb-Scargle periodogram analysis on* ***incubation recesses****, extracted from wildlife camera pictures. Level of significance is set to P < 0.05. One goose did not show a significant diel peak period.*

| **Goose ID** | **Peak period ultradian (h)** | **Peak power ultradian** | **Significance threshold ultradian** | **Peak period diel (h)** | **Peak power diel** | **Significance threshold diel** |
|---|---|---|---|---|---|---|
| CA41113 | 3.63 | 0.02 | 0.005 | 23.70 | 0.03 | 0.003 |
| CA41238 | 2.78 | 0.02 | 0.004 | 24.22 | 0.03 | 0.003 |
| CA41400 | 6.48 | 0.02 | 0.004 | 22.78 | 0.02 | 0.003 |
| CA44228 | 3.49 | 0.01 | 0.004 | 24.37 | 0.02 | 0.003 |
| CA45632 | 3.60 | 0.03 | 0.005 | 23.74 | 0.05 | 0.004 |
| CA45701 | 3.62 | 0.02 | 0.004 | 24.01 | 0.02 | 0.003 |
| CA45736 | 1.76 | 0.03 | 0.006 | | | |
| CA45744 | 3.43 | 0.02 | 0.005 | 24.32 | 0.02 | 0.004 |
| CA45891 | 2.27 | 0.03 | 0.005 | 23.81 | 0.04 | 0.003 |
| CA46848 | 1.94 | 0.02 | 0.005 | 24.64 | 0.02 | 0.003 |
| CA45997 | 2.27 | 0.02 | 0.005 | 24.67 | 0.02 | 0.004 |

*Table 2: Results of the Lomb-Scargle periodogram analysis on* ***sleep posture****, extracted from wildlife camera pictures. Level of significance is set to* $P < 0.05$*. Two geese did not show significant ultradian peak periods.*

| **Goose ID** | **Peak period ultradian (h)** | **Peak power ultradian** | **Significance threshold ultradian** | **Peak period diel (h)** | **Peak power diel** | **Significance threshold diel** |
|---|---|---|---|---|---|---|
| CA41113 | 3.63 | 0.02 | 0.004 | 23.75 | 0.05 | 0.003 |
| CA41238 | 2.78 | 0.01 | 0.003 | 23.07 | 0.02 | 0.002 |
| CA41400 | 6.45 | 0.01 | 0.004 | 23.87 | 0.03 | 0.003 |
| CA44228 | 12.37 | 0.02 | 0.004 | 23.82 | 0.02 | 0.003 |
| CA45632 | 5.98 | 0.03 | 0.005 | 26.46 | 0.03 | 0.003 |
| CA45701 | 12.45 | 0.02 | 0.004 | 23.87 | 0.03 | 0.003 |
| CA45736 | 4.64 | 0.02 | 0.006 | 26.66 | 0.03 | 0.004 |
| CA45744 | 9.96 | 0.01 | 0.004 | 30.87 | 0.02 | 0.003 |
| CA45891 | | | | 23.12 | 0.07 | 0.003 |
| CA46848 | | | | 23.87 | 0.04 | 0.003 |
| CA45997 | 15.37 | 0.03 | 0.004 | 25.21 | 0.05 | 0.003 |

#### *Seasonal rhythmicity in activity*

*Table 3: The 95% confidence set of models investigating differences in* ***ultradian peak period in hours in activity behaviour during three breeding stages*** *(nesting, gosling, moult periods) within a cumulative Akaike weight of ≥ 0.95 from the top model. The '+' indicates which variables are present in the different models, delta gives the difference in AICc between the models, and weight represents the relative probability that a certain model is best compared with a set of models, based on the data and taking model complexity into account.*

| **Intercept** | **sex** | **breeding stage** | **year** | **sex : breeding stage** | **df** | **logLik** | **AICc** | **delta** | **weight** |
|---|---|---|---|---|---|---|---|---|---|
| 5.36 | | + | | | 5 | -73.14 | 159.01 | 0.00 | 0.39 |
| 5.36 | | + | + | | 6 | -71.55 | 159.11 | 0.09 | 0.37 |
| 6.28 | + | + | | | 6 | -72.55 | 161.10 | 2.08 | 0.14 |

*Table 4: The 95% confidence set of models investigating differences in* ***diel peak period in hours in activity behaviour during three breeding stages*** *(nesting, gosling, moult periods) within a cumulative Akaike weight of ≥ 0.95 from the top model. The '+' indicates which variables are present in the different models, delta gives the difference in AICc between the models, and weight represents the relative probability that a certain model is best compared with a set of models, based on the data and taking model complexity into account.*

| Intercept | sex | breeding stage | year | sex : breeding stage | df | logLik | AICc | delta | weight |
|---|---|---|---|---|---|---|---|---|---|
| 23.61 | | | | | 3 | -74.04 | 154.89 | 0.00 | 0.49 |
| 23.82 | | | + | | 4 | -73.72 | 156.82 | 1.93 | 0.19 |
| 23.35 | + | | | | 4 | -73.82 | 157.02 | 2.13 | 0.17 |
| 23.55 | + | | + | | 5 | -73.48 | 159.10 | 4.21 | 0.06 |

#### ***Rhythmicity in corticosterone***

*Table 5: The 95% confidence set of linear fixed-effects models investigating rhythmicity in log-transformed* ***corticosterone metabolite concentrations*** *over the course of the day within a cumulative Akaike weight of ≥ 0.95 from the top model. The '+' indicates which variables are present in the different models, delta gives the difference in AICc between the models, and weight represents the relative probability that a certain model is best compared with a set of models, based on the data and taking model complexity into account.*

| Intercept | cosine | sine | sex | year | day of the year | df | logLik | AICc | Delta | weight |
|---|---|---|---|---|---|---|---|---|---|---|
| 4.99 | 0.10 | 0.06 | | + | -0.007 | 7 | -708.88 | 1431.93 | 0.00 | 0.31 |
| 4.89 | 0.11 | | | + | -0.007 | 6 | -710.02 | 1432.16 | 0.23 | 0.28 |
| 5.03 | 0.10 | 0.06 | + | + | -0.007 | 8 | -708.51 | 1433.23 | 1.30 | 0.16 |
| 4.93 | 0.10 | | + | + | -0.007 | 7 | -709.66 | 1433.50 | 1.56 | 0.14 |
| 4.85 | | 0.06 | | + | -0.007 | 6 | -712.57 | 1437.26 | 5.33 | 0.02 |
| 4.75 | | | | + | -0.006 | 5 | -713.87 | 1437.82 | 5.89 | 0.02 |
| 4.90 | | 0.06 | + | + | -0.007 | 7 | -712.16 | 1438.48 | 6.55 | 0.01 |
| 4.35 | 0.10 | | | | -0.005 | 5 | -714.46 | 1439.01 | 7.07 | 0.01 |

*Table 6: The 95% confidence set of linear fixed-effects models investigating rhythmicity in a* ***subset of log-transformed corticosterone metabolite concentrations*** *over the course of the day, i.e. excluding samples with concentrations of >200 ng CORTm/g droppings (n=7), within a cumulative Akaike weight of ≥ 0.95 from the top model. The '+' indicates which variables are present in the different models, delta gives the difference in AICc between the models, and weight represents the relative probability that a certain model is best compared with a set of models, based on the data and taking model complexity into account.*

| **Intercept** | **cosine** | **sine** | **sex** | **year** | **day of the year** | **df** | **logLik** | **AICc** | **delta** | **weight** |
|---|---|---|---|---|---|---|---|---|---|---|
| 3.51 | 0.07 | | | + | | 5 | -658.32 | 1326.74 | 0.00 | 0.18 |
| 4.05 | 0.08 | | | + | -0.003 | 6 | -657.50 | 1327.13 | 0.39 | 0.15 |
| 3.55 | 0.07 | | + | + | | 6 | -657.83 | 1327.78 | 1.05 | 0.10 |
| 4.09 | 0.07 | | + | + | -0.003 | 7 | -657.00 | 1328.17 | 1.43 | 0.09 |
| 3.49 | | | | + | | 4 | -660.25 | 1328.57 | 1.83 | 0.07 |
| 3.51 | 0.07 | 0.01 | | + | | 6 | -658.25 | 1328.63 | 1.89 | 0.07 |
| 4.09 | 0.07 | 0.02 | | + | -0.003 | 7 | -657.33 | 1328.83 | 2.09 | 0.06 |
| 3.54 | | | + | + | | 5 | -659.71 | 1329.52 | 2.78 | 0.04 |
| 3.93 | | | | + | -0.002 | 5 | -659.71 | 1329.52 | 2.78 | 0.04 |
| 3.55 | 0.07 | 0.01 | + | + | | 7 | -657.75 | 1329.67 | 2.94 | 0.04 |
| 4.13397 | 0.07 | 0.02 | + | + | -0.003 | 8 | -656.82 | 1329.86 | 3.12 | 0.04 |
| 3.49 | | 0.02 | | + | | 5 | -660.14 | 1330.37 | 3.64 | 0.03 |
| 3.97 | | | + | + | -0.002 | 6 | -659.16 | 1330.46 | 3.72 | 0.03 |

*Table 7: Intercept and coefficient estimates from the full model investigating rhythmicity in a* ***subset of log-transformed corticosterone metabolite concentrations****, i.e. excluding samples with concentrations of >200 ng CORTm/g droppings with corresponding 85% CIs, and if the variable was selected in the top AIC model.*

| **Variable** | **estimate** | **SE** | **Lower** | **upper** | **AIC top** |
|---|---|---|---|---|---|
| Intercept | 4.13 | 0.43 | 3.52 | 4.75 | |
| cosine | 0.073 | 0.04 | 0.02 | 0.13 | Yes |
| sine | 0.02 | 0.04 | -0.03 | 0.07 | No |
| sex - male | -0.09 | 0.09 | -0.22 | 0.04 | No |
| year - 2021 | -0.22 | 0.06 | -0.31 | -0.13 | Yes |
| year day | -0.003 | 0.002 | -0.006 | 0.0001 | No |

##### *Seasonal rhythmicity in activity*

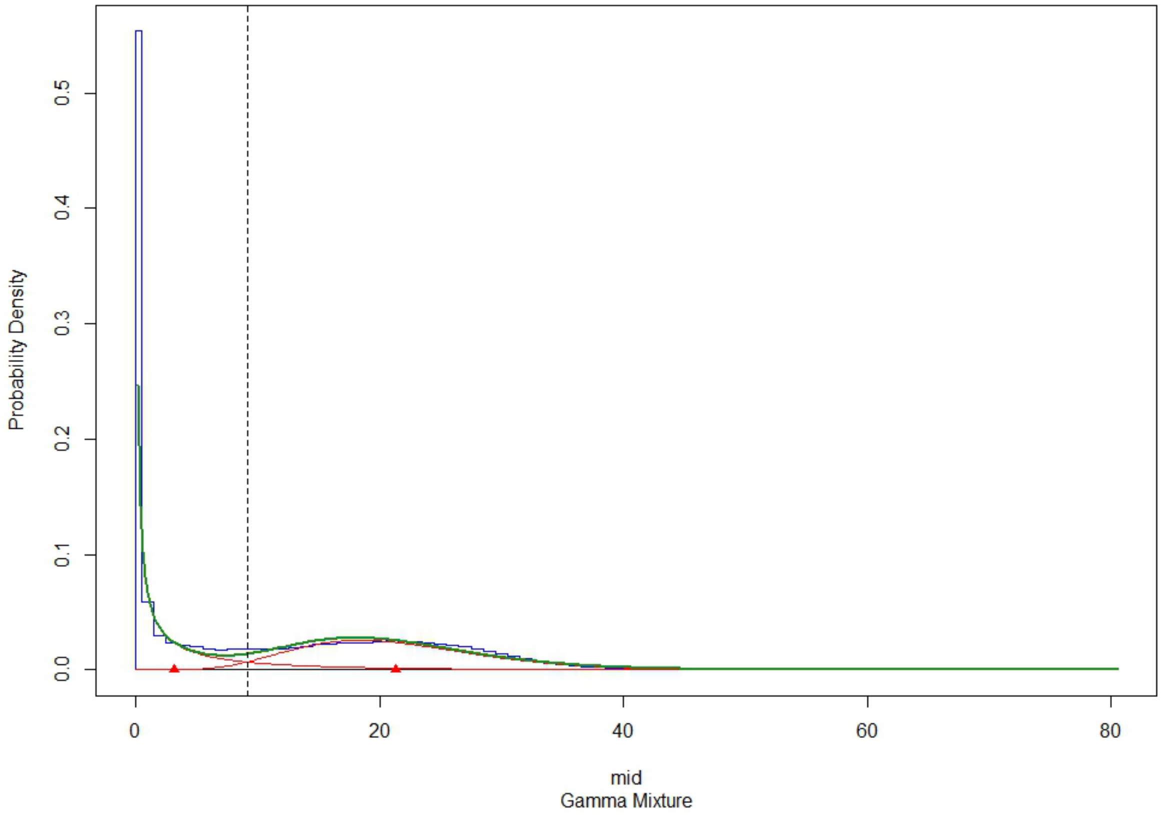


*Figure 1: Probability density histogram for* ***vectoral dynamic body acceleration (VeDBA).*** *The blue line shows the probability density histogram, the green lines show the corresponding probability density functions and the red lines depict the fitted gamma distributions. The red triangles give the mid points of the gamma distributions for inactive and active behaviour. The dashed vertical line is the intersection point between the distributions and as such gives the threshold (9.15) used to distinguish VeDBA values indicating active and inactive behaviour.*

#### *Incubation behaviour*

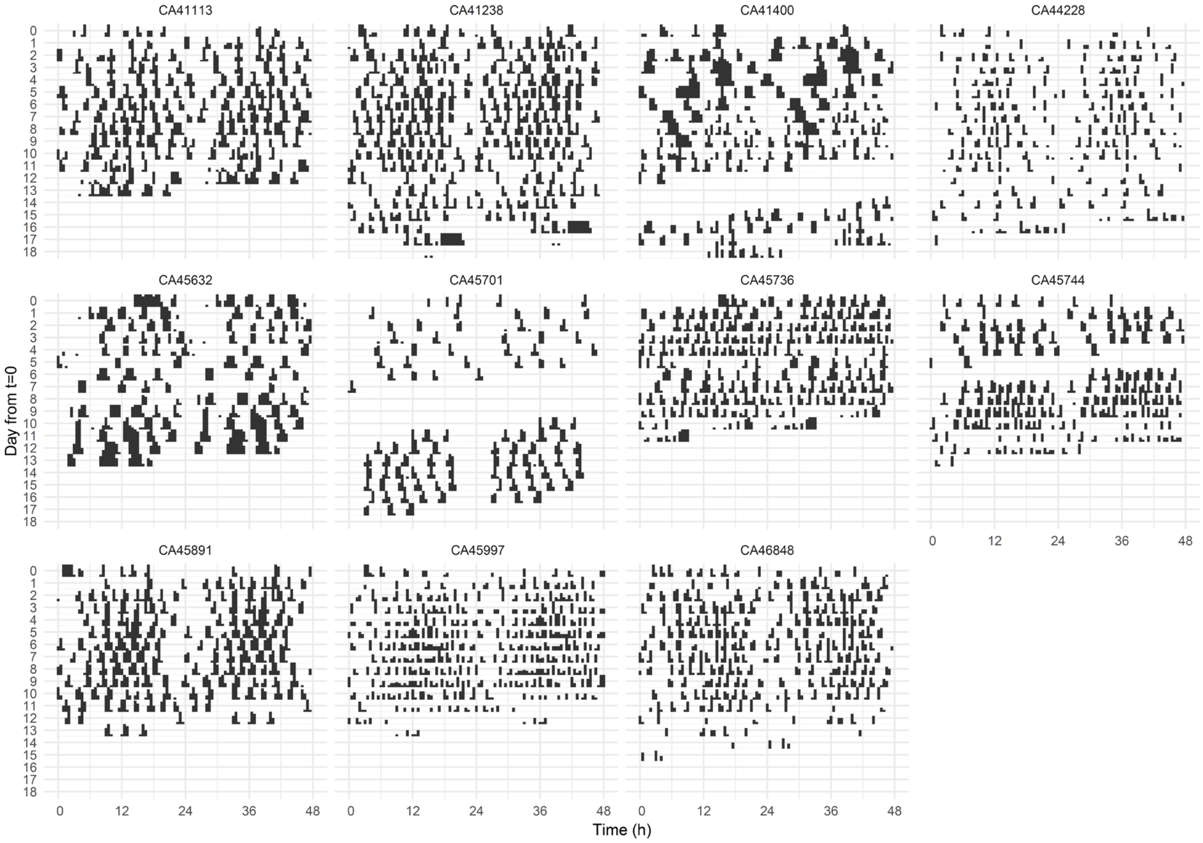


*Figure 2: Double-plotted actograms of* ***incubation recesses*** *for all individual females. Day t = 0 is measured from when the camera was set up near the nest. All but one female CA45701 (second panel middle row), whose nest failed for unknown reasons, had successful nests. In three females data are incomplete due to camera malfunction: in CA41400 data for two days are missing, in CA45701 data for three days are missing, and in CA45744 data for one day are missing.*

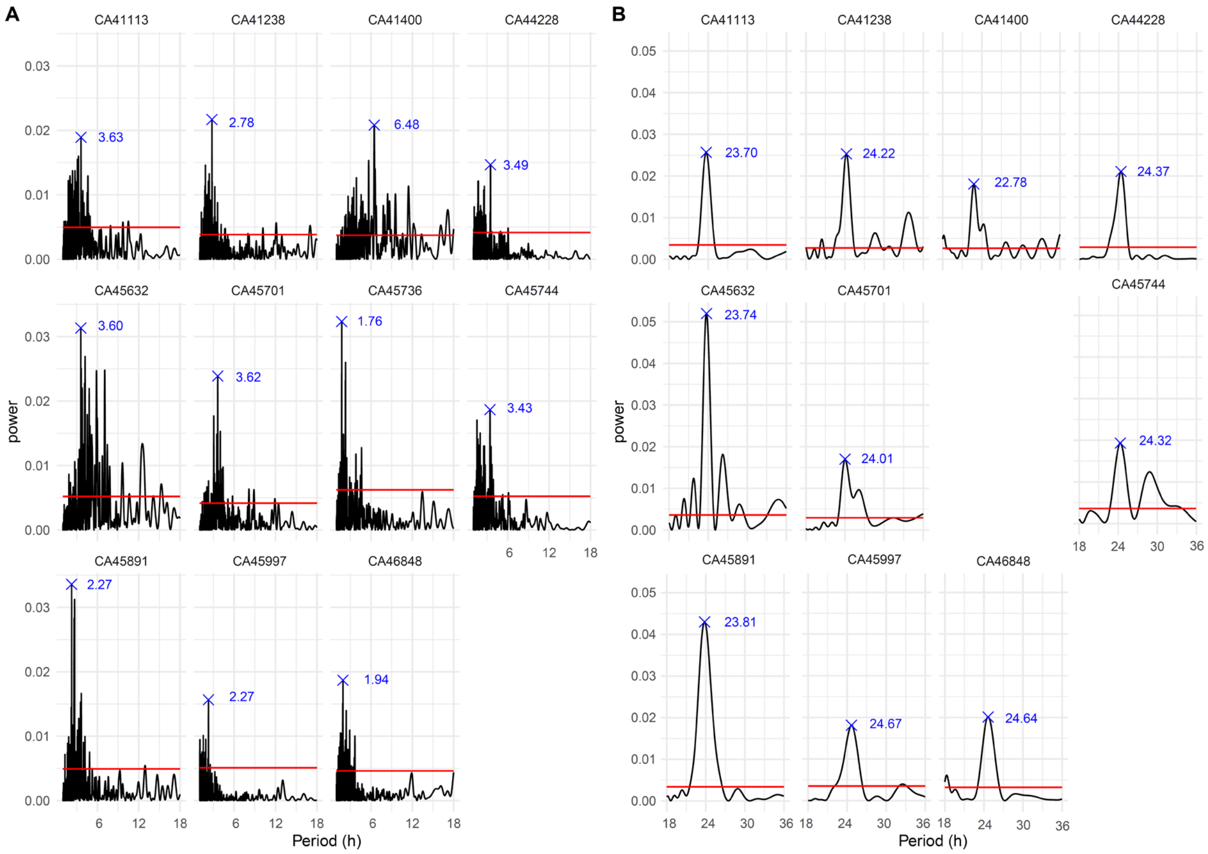


*Figure 3: Lomb-Scargle periodograms in **incubation recesses** for all individual geese. The analysis identified periodicity between A) 1 and 18 hours (to focus on ultradian rhythmicity) and B) between 18 and 36 hours (to focus on diel rhythmicity). Peaks above the significance threshold of P < 0.05 (red line) are shown in blue. With the exception of CA45736, who only showed ultradian rhythmicity (third panel middle row), all geese showed both ultradian and diel rhythmicity. All geese had successful nests except for CA45701 whose nest failed because of unknown reasons.*

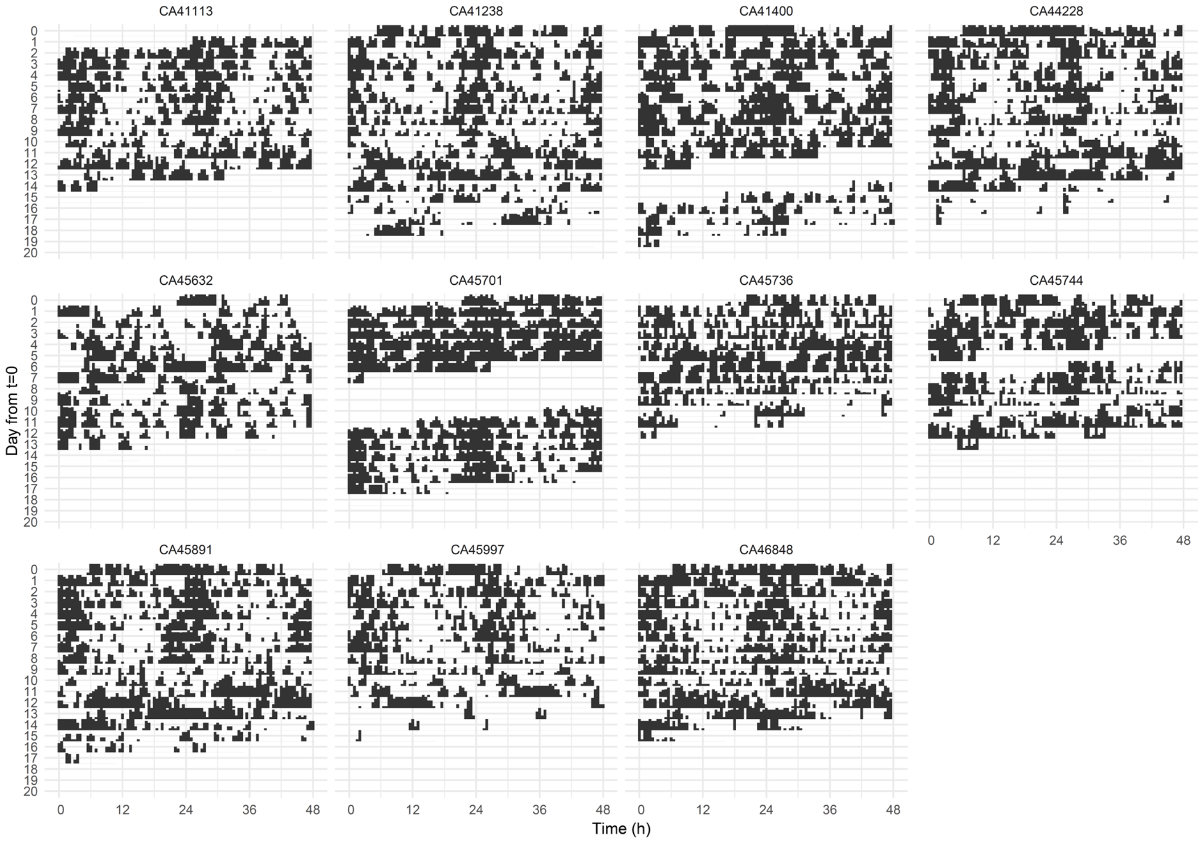


*Figure 4: Double-plotted actograms of* ***sleep posture*** *for all individual females. Day t = 0 is measured from when the camera was set up near the nest. All but one female CA45701 (second panel middle row), whose nest failed for unknown reasons, had successful nests. In three females, data are incomplete due to camera malfunction: in CA41400 data for two days are missing, in CA45701 data for three days are missing, and in CA45744 data for one day are missing.*

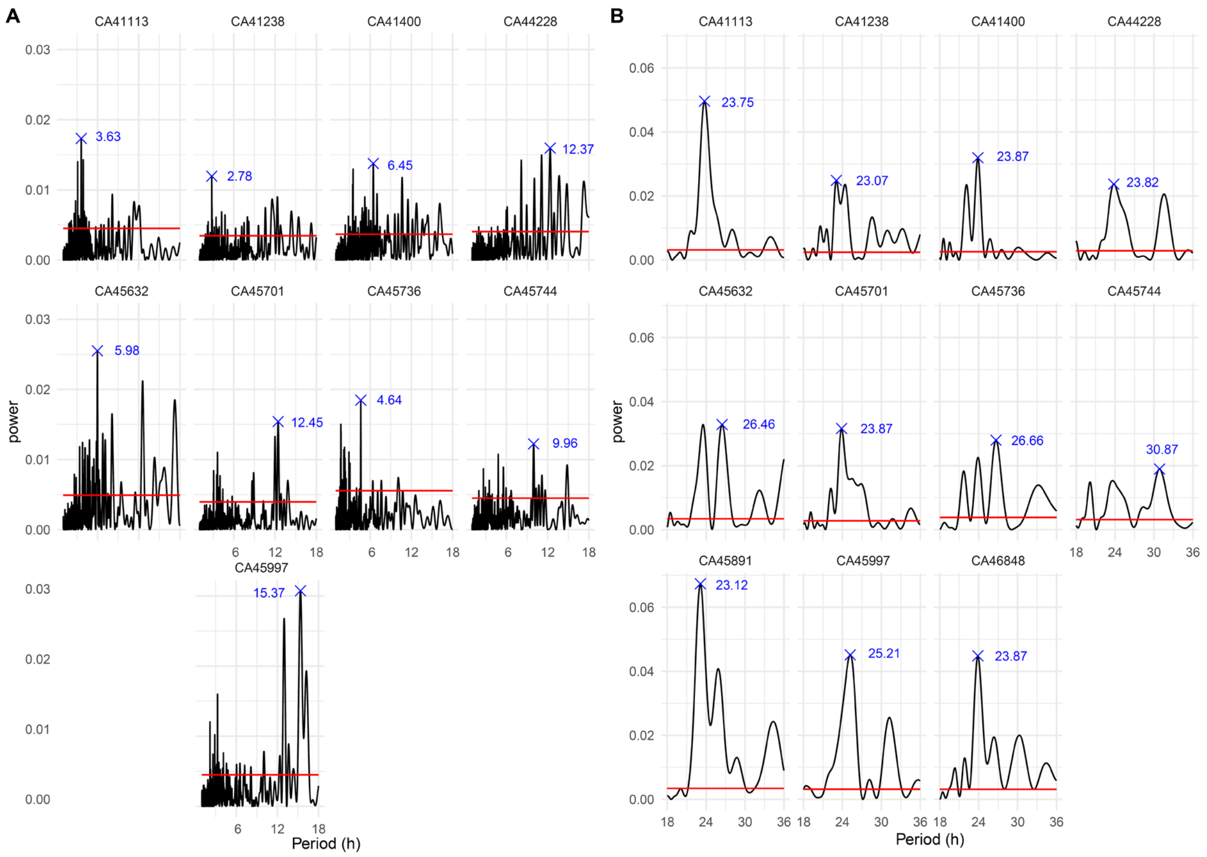


*Figure 5: Lomb-Scargle periodograms of* ***sleep posture*** *for all individual geese. The analysis identified periodicity between (A) 1 and 18 hours (to focus on ultradian rhythmicity) and (B) between 18 and 36 hours (to focus on diel rhythmicity). Peaks above the significance threshold of* $P < 0.05$ *(red line) are shown in blue. With two exceptions, CA45891and CA46848, who only showed diel rhythmicity (first and third panel bottom row) geese showed both ultradian and diel rhythmicity. All geese had successful nests except for CA45701 whose nest failed because of unknown reasons.*

***Seasonal rhythmicity in activity***

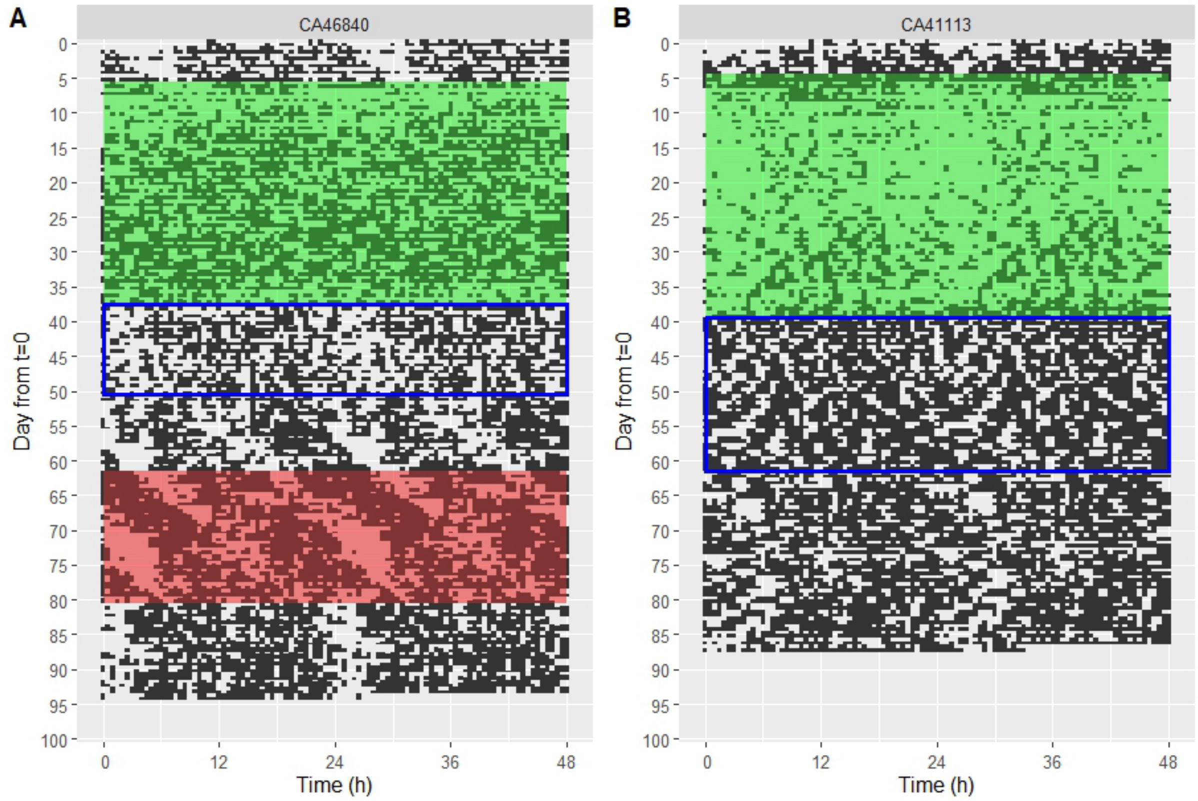


*Figure 6: Examples of rhythmicity in* ***activity over the course of the season*** *in two geese: (A) Double-plotted actogram of a male goose (ID CA46840) with ultradian and circadian rhythmicity during nesting (in green) and when it was observed with goslings (blue rectangle), but only ultradian rhythmicity during moult (in red), (B) a female goose (ID CA41113) with circadian rhythmicity during nesting and ultradian and circadian rhythmicity when it was observed with goslings (blue rectangle). This female was not observed during moult. In double-plotted actograms, the x-axis displays two consecutive days, and these consecutive days are also shown from top to bottom on the y-axis. Activity is shown in black, while transparency indicates inactivity.*
